# Matrix and analysis metadata standards (MAMS) to facilitate harmonization and reproducibility of single-cell data

**DOI:** 10.1101/2023.03.06.531314

**Authors:** Yichen Wang, Irzam Sarfraz, Wei Kheng Teh, Artem Sokolov, Brian R. Herb, Heather H. Creasy, Isaac Virshup, Ruben Dries, Kylee Degatano, Anup Mahurkar, Daniel J Schnell, Pedro Madrigal, Jason Hilton, Nils Gehlenborg, Timothy Tickle, Joshua D. Campbell

**Affiliations:** Department of Medicine, Boston University School of Medicine, Boston, MA, USA; European Bioinformatics Institute, European Molecular Biology Laboratory, Hinxton, Cambridgeshire, UK; Laboratory of Systems Pharmacology, Harvard Medical School, Boston, MA, USA; Institute for Genome Sciences, University of Maryland School of Medicine, Baltimore, MD, USA; Department of Computational Health, Helmholtz Munich, Oberschleißheim, Germany; Data Sciences Platform, Broad Institute, Cambridge, MA, USA; Biomedical Informatics, Cincinnati Children’s Hospital Medical Center, Cincinnati, OH, USA; Department of Genetics, Stanford University, Stanford, CA, USA; Biomedical Informatics, Harvard Medical School, Boston, MA, USA

## Abstract

A large number of genomic and imaging datasets are being produced by consortia that seek to characterize healthy and disease tissues at single-cell resolution. While much effort has been devoted to capturing information related to biospecimen information and experimental procedures, the metadata standards that describe data matrices and the analysis workflows that produced them are relatively lacking. Detailed metadata schema related to data analysis are needed to facilitate sharing and interoperability across groups and to promote data provenance for reproducibility. To address this need, we developed the Matrix and Analysis Metadata Standards (MAMS) to serve as a resource for data coordinating centers and tool developers. We first curated several simple and complex “use cases” to characterize the types of feature-observation matrices (FOMs), annotations, and analysis metadata produced in different workflows. Based on these use cases, metadata fields were defined to describe the data contained within each matrix including those related to processing, modality, and subsets. Suggested terms were created for the majority of fields to aid in harmonization of metadata terms across groups. Additional provenance metadata fields were also defined to describe the software and workflows that produced each FOM. Finally, we developed a simple list-like schema that can be used to store MAMS information and implemented in multiple formats. Overall, MAMS can be used as a guide to harmonize analysis-related metadata which will ultimately facilitate integration of datasets across tools and consortia. MAMS specifications, use cases, and examples can be found at https://github.com/single-cell-mams/mams/.

## INTRODUCTION

The past two decades have seen a rapid expansion of high throughput genomic and imaging technologies that have revolutionized the ability of researchers to capture the molecular and histological characteristics of biological samples. For example, assays such as single-cell RNA-seq can capture the states of individual cells within heterogeneous and complex tissues. Several major consortia have been funded to utilize single cell assays to create cellular atlases of healthy and disease tissues^1–7^. These groups are generating large datasets that contain multi-modal single-cell data collected from longitudinally- and spatially-related biological specimens from different organs across different conditions. The majority of datasets are designed to answer a specific set of questions within a particular biological or clinical context. Other data centers such as NIH Gene Expression Omnibus (GEO) and ArrayExpress/BioStudies databases are focused on systematic storage, cataloging, and retrieval of primary data^8,9^. As more data becomes available, the ability to combine and integrate datasets from different settings is becoming increasingly desirable. Three major roadblocks to combining and integrating datasets are that 1) the metadata related to clinical, biospecimen, and experimental parameters is not captured or harmonized across groups^10^; 2) the data is stored in a wide variety of file formats or programming language-specific libraries, classes, or data structures; and 3) the metadata about the matrix and the corresponding analysis that produced or utilized the matrix is not well standardized. While significant effort has been dedicated to defining metadata standards for experimental parameters^10^ and new file formats are under active development^11^, the third area remains largely unaddressed.

While a wide range of experimental protocols and platforms are available to generate molecular and histological data, an important commonality across these technologies is that they often produce a matrix of features that are measured in a set of observations. These feature and observation matrices (FOMs) are foundational for storing raw data from molecular assays (e.g. raw counts) and derived data from downstream analytical tools (e.g. normalized matrix, reduced dimensional matrix). A variety of file formats are used to store FOMs on file systems in different representations. For example, Tab Separated Value (tsv/txt) files can be used to store dense matrices while Market Exchange (.mtx) files can be used to efficiently store sparse matrices. Although platform-independent, these formats do not readily capture relationships between matrices and annotations for features and observations. Several packages also exist that can capture relationships between matrices and annotations include AnnData and MUON in Python^12,13^, the Seurat object in R^14,15^, and the SingleCellExperiment and related packages in R/Bioconductor^16–18^. In contrast to the simple flat file formats, these objects can capture complex relationships between FOMs as well as annotation data produced during the analysis such as quality control (QC metrics) and cluster labels. However, each package may store different sets of data or label the same type of data differently. Even if different datasets are stored the same format or type of object, harmonization of datasets still may require substantial manual curation before they can be combined and integrated.

Lastly, a major goal of high-quality analysis workflows is to promote reproducibility by storing information related to provenance such as software version, function calls, and selected parameters that produced the matrix or annotation. However, there is a high degree of variability in which different analytical tools and software packages capture this information. Even if the data was produced by a workflow captured in a Docker container or versioned in GitHub, this information will often be lost when converting the data between formats or transferring between tools. Thus, there is a need to develop metadata standards for FOMs related to provenance to ensure this information can be readily captured and maintained throughout the dataset-specific analysis and during integration with other datasets.

In order to facilitate sharing and interoperability across groups and technologies as well as to promote reproducibility related to data provenance, a detailed metadata schema describing the characteristics of FOMs can be used to serve as a standard for the community. Therefore, we developed the **Matrix and Analysis Metadata Standards (MAMS)** to capture the relevant information about the data matrices and annotations that are produced during common and complex analysis workflows for single-cell data. MAMS defines fields that describe what type of data is contained within a matrix, relationships between matrices, and provenance related to the tool or algorithm that created the matrix. In contrast to the existing standards, MAMS does not largely focus on information related to sample preparation including biospecimen and clinical metadata or metadata related to experimental protocols. These standards will serve as a roadmap for tool developers and data curators to ensure that their systems have the capability to store and retrieve relevant information needed for integration. All of the metadata fields are independent of the platform, programming language, and specific tool and thus can be used to support efforts to harmonize data across consortia.

## RESULTS

### Overview of matrix classes

Several popular libraries and software packages offer convenient interfaces for storing and retrieving data matrices and their associated annotations. The majority of these tools employ similar schemas that organize different classes of data matrices in an intuitive framework with a common interface (**Figure 1**). In general, we refer to a feature and observation matrix (FOM) as a class of data matrix that contains measurements of features across biological entities. Examples of features include genes, genomic regions, peaks, transcripts, proteins, antibodies derived tags, signal intensities, cell type counts or morphology categories. Examples of observations include cells, cell pools, beads, spots, subcellular regions, and regions of interest (ROIs). Measurements for single-cell data may include transcript counts, protein abundances, signal intensities and velocity estimates. FOMs that contain raw, normalized, transformed, or standardized biological data are commonly referred to as “assays” or “layers”. In the MAMS nomenclature, FOMs can also contain reduced dimensional objects such as principal components from PCA or 2-D embedding from tSNE or UMAPs which are derived from the original biological data matrices. We note that although the term “matrix” is used in the acronym, FOMs may also be data frames which can contain mixed data types (e.g. continuous and categorical morphological features), vectors (e.g. a matrix of with 1 dimension), and multidimensional arrays (e.g. a matrix with more than 2 dimensions also known as a tensor).

**Figure 1.**
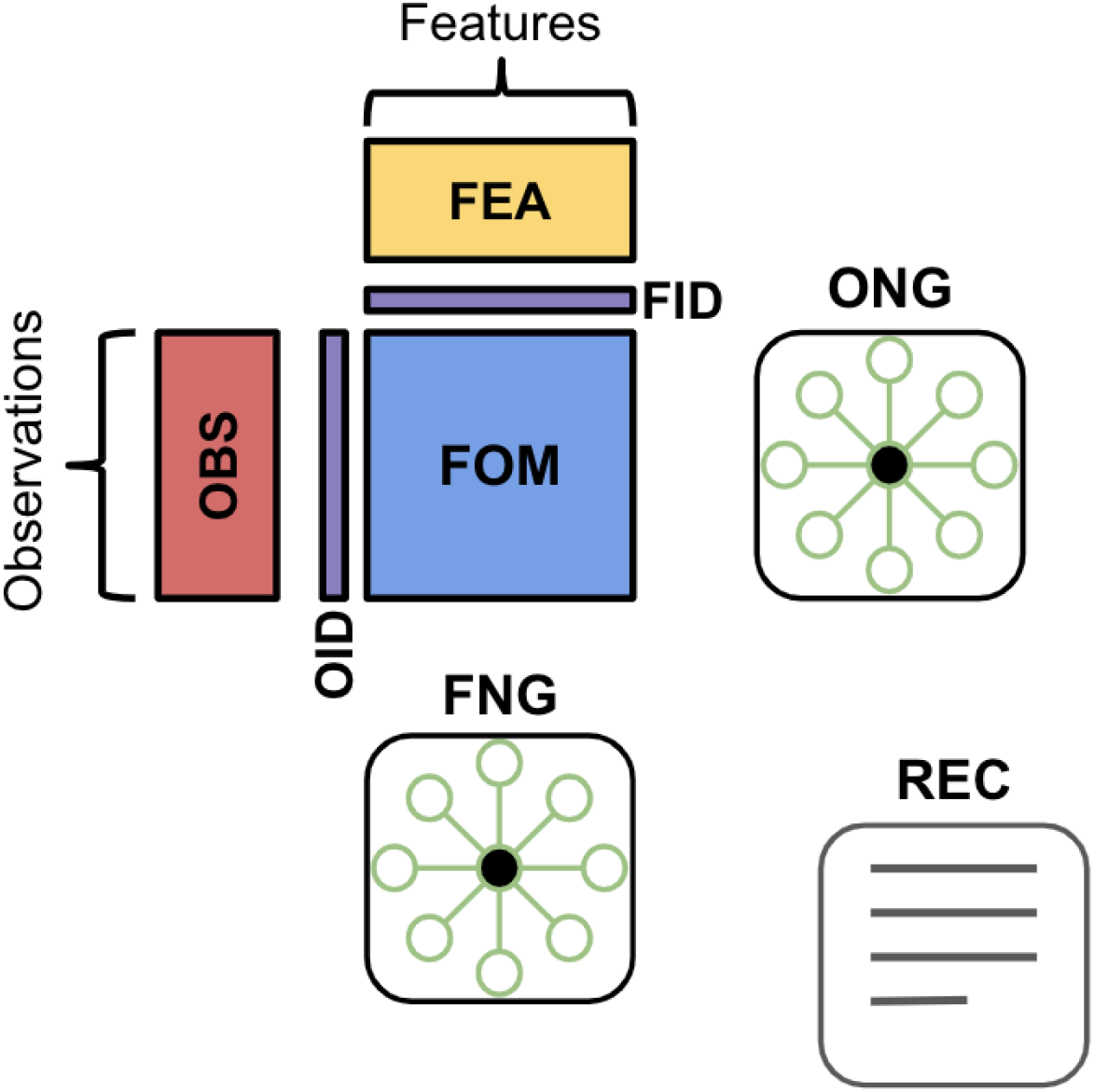
Overview of matrix classes included in MAMS. Feature and observation matrices (FOMs) contain biological data at different stages of processing including reduced dimensional representations. Feature annotation matrices (FEA) and Observation annotation matrices (OBS) store annotations such as additional IDs or labels, quality control metrics, and cluster labels. The Observation Neighborhood Graph (ONG) and Feature Neighborhood Graph (FNG) classes store information related to the correlation, similarity, or distance between pairs of observations or features, respectively. The Observation ID (OID) and Feature ID classes are used to store unique identifiers for individual observations and features, respectively. The Record (REC) class is a special set of fields for storing information related to data and tool provenance.

Beyond the central data matrices, other classes of matrices are used to store identifier (IDs) information, annotations, and graphs that are generated during analysis. Annotations and metadata about the features (FEA) or observations (OBS) are stored in separate data frames with the same dimensions as the parent FOM. Annotations for features can include information about the gene such as IDs, reference genome, genomic location, biotype, and variability metrics. Annotations for observations can include sample-level demographics, cell-level identifiers (e.g. barcode), quality control metrics (e.g. total number of features detected), or analysis output (e.g. cluster labels, trajectory scores). IDs are used to uniquely identify and index observations and features within a dataset. The Observation ID (OID) class contains a character vector or combination of character vectors used to denote the unique ID of each observation while the Feature ID (FID) class contains a vector or combination of character vectors used to denote the unique ID of each feature. Observation neighborhood graphs (ONGs) and feature neighborhood graphs (FNGs) are adjacency matrices that can be used to store the correlation, similarity, or distance between pairs of observations and features, respectively. In the MAMS schema, each class of matrix will have a corresponding set of metadata fields that describe the information contained within the matrix. The Record (REC) is a special class for storing information related to the provenance of the data analysis tool and command used to create the matrix.

### Curated analysis workflows for single cell data

In order to ensure that the metadata standards are able to capture complex information and robust to different real-world scenarios, we first curated the matrices produced by different types of analysis workflows for singlecell data using the vignettes from popular tools and packages as a template^14,19,20^. The simplest workflow starts with a matrix of UMI-correct counts produced by a microfluidics device where the observations are droplets denoted with a unique barcode, the features are genes, and the measurements are the number of mRNA transcripts detected for each gene in each cell (**Figure 2**). This type of matrix is produced by aligning sequencing reads to a reference genome, counting the number of reads that map to each gene loci, and then performing a correction for Unique Molecular Indices (UMIs). The next step is to identify and filter the observations of the matrix to remove empty droplets (i.e. droplets without a true cell). The observations in this filtered matrix can be filtered again based on other quality control metrics such as total number of UMIs detected, number of features detected, percentage of mitochondrial counts, percentage of ambient RNA, or droplets likely containing multiple cells. After generating a “clean” matrix of observations, the raw counts are generally normalized by correcting for library size (e.g. correcting for the total number of counts) and applying a log2 transformation. Next, features are standardized across observations (e.g. z-scoring each gene to have a mean of 0 and a standard deviation of 1) and a subset of highly variable features are chosen for downstream analysis. This matrix is used as input into Principal Component Analysis (PCA) to produce a reduced dimensional matrix, where the features are now principal components (PCs) instead of individual genes. Optionally, a subset of PCs may be selected based on percentage of variability or other statistical metrics.

**Figure 2.**
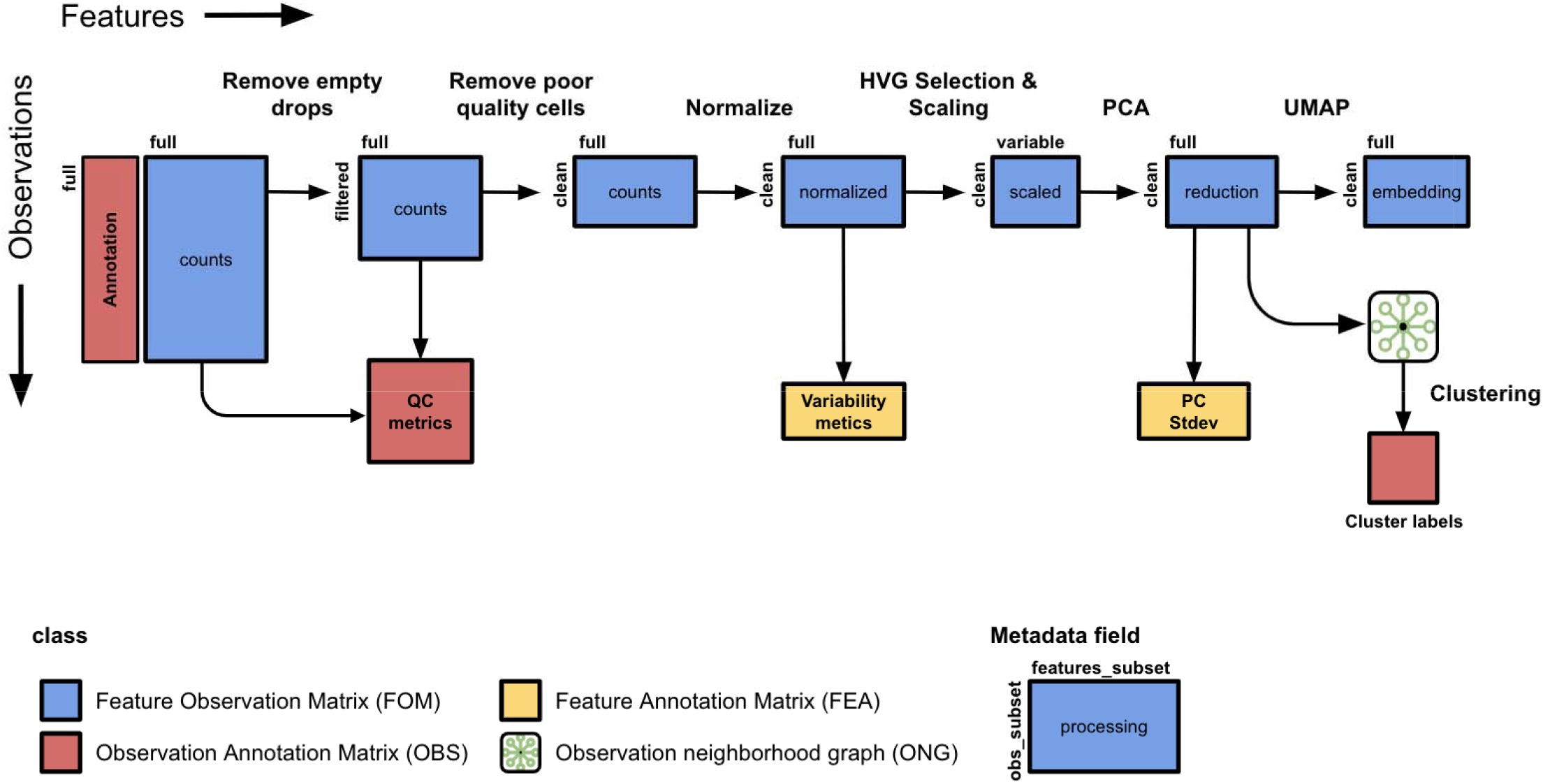
Matrices produced during a simple analysis workflow for single cell RNA-seq data. Several steps are often performed in analysis workflows for scRNA-seq data generated with high-throughput devices. The observations are filtered to exclude empty droplets and poor quality cells. Quality control metrics can be stored in an OBS annotation data frame. Preprocessing of the data matrix includes steps for normalization and standardization of features (e.g. z-scoring). From the scaled data, a subset of highly variable genes are used as input into Principal Component Analysis (PCA). The reduced dimensional space of the PCA is used as input into 2D embedding tools such as tSNE and UMAP as well as clustering algorithms such as k-means and Leiden.

**Figure 3.**
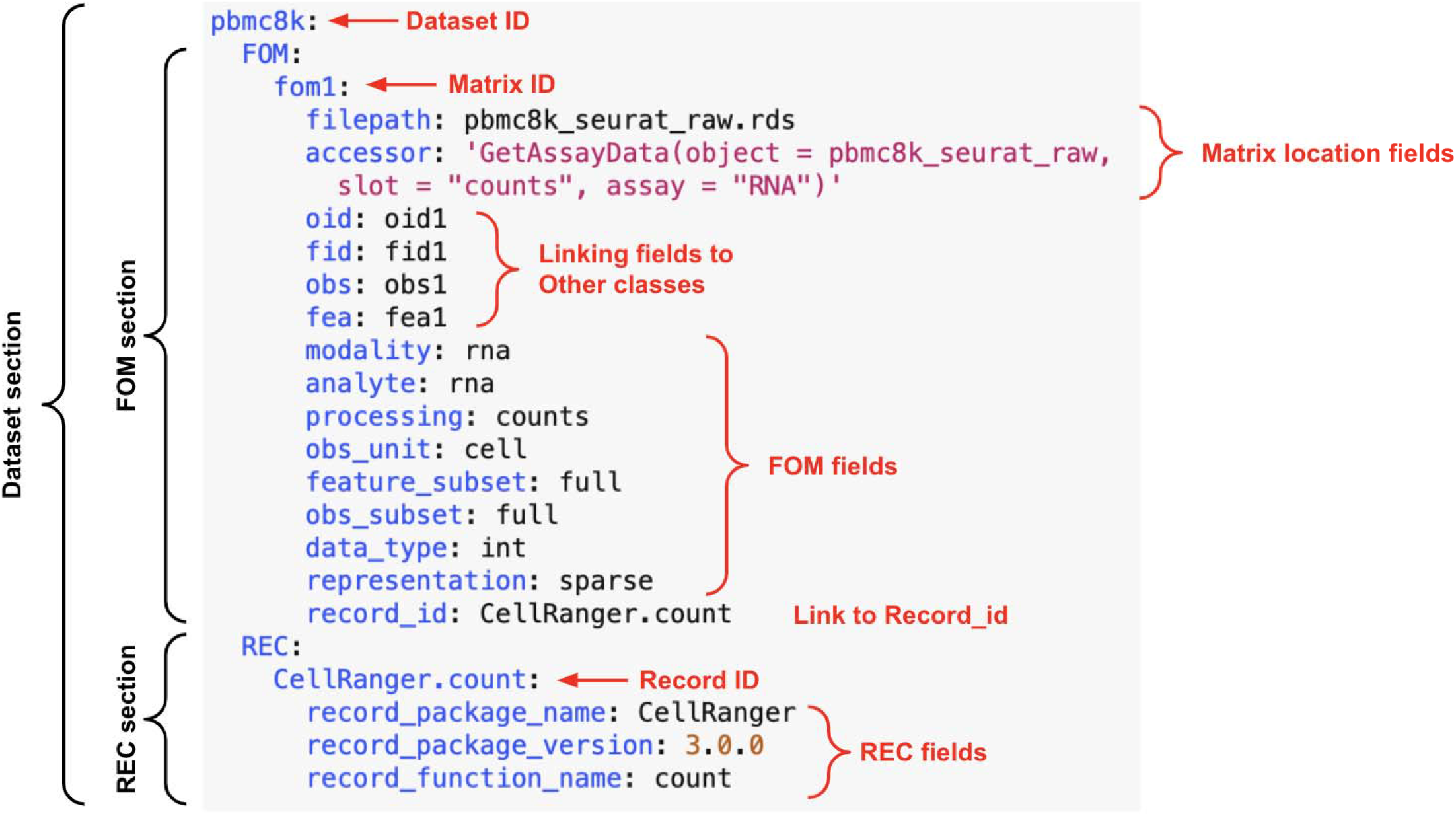
Example of MAMS list format. As the ability to implement and store matrix and analysis related metadata is variable across software platforms and data objects, we created a simple list-like structure to capture relevant MAMS fields for each matrix. This structure can be stored in configuration file formats like JSON and YAML or in general metadata or unstructured slots within data objects. Each dataset will have its own entry within the list and each class of matrix has an entry within the list for each dataset. Each matrix is denoted with a unique ID and MAMS fields are denoted with key-value pairs under each matrix. The additional fields specified within this implementation including ***filepath*** and ***accessor*** can be used to point to matrices stored in any flat file format or within a data object.

Neighborhood and distance graphs are produced between observations using the PCA matrix as input. These distances are used as input into algorithms such as tSNE and UMAP that produce another matrix of 2-D embeddings useful for visualization. The cell graphs can also be used as input into clustering algorithms such as Leiden or K-means or trajectory inference tools. Annotations such as cluster labels and trajectory scores are stored in the corresponding OBS data frame with the same number of observations. A robust analysis metadata standard should be able to capture information about the processing of the measurements with each FOM as well as the features and observations included in each FOM.

While the simple workflow contains analysis of a single sample, other scenarios may require more complex workflows with additional types of operations that produce different matrices. We also annotated scenarios that include analysis of datasets with multiple modalities (**Supplementary Figure 1**), datasets with multiple samples that require integration or batch correction (**Supplementary Figure 2**), analysis with a biological subset of data (**Supplementary Figure 3**), and analysis with FOMs derived from imaging data (**Supplementary Figure 4**). Multiple modalities can be measured on individual cells in addition to mRNA transcript counts. For example, scATAC-seq assays can measure open chromatin profiles and CITE-seq assays can measure epitopes for cell surface proteins. Multimodal workflows may apply similar sets of procedures to each data type to normalize data, perform dimensionality reduction, and generate modalityspecific cluster labels. To perform integrative analysis, matrices from different modalities can be integrated at the “matrix” level, reduced dimensional level, or on the graph level^21^ and a third set of cluster labels can be derived using the combined dataset. A robust analysis metadata standard should be able to capture information about the biological analyte and modality captured with each FOM.

Other single-cell datasets may have observations from multiple donors or multiple regions per donor. Several batch correction and integration tools have been developed to remove unwanted variation between different samples and project the shared variation into a common low dimensional representation which can be used in subsequent graph generation and clustering steps (**Supplementary Figure 2**). After initial analysis using all observations that passed quality control, additional analyses on subsets of biologically may be desired. For example, initial clustering with single cell data may be used to define broad cell types such as epithelial, stromal, and immune cells. The cells from each broader cell type may be subsetted and re-analyzed using a similar workflow to produce novel matrices and annotations (**Supplementary Figure 3**). A robust analysis metadata standard should be able to capture information about the batch corrections and biological subsets.

Lastly, we curated workflows where the underlying data was derived from highly-multiplexed imaging technologies, such as cyclic immunofluorescence (CyCIF), CODEX, and MIBI which measure multiple biological features on the same tissue slide (**Supplementary Figure 4**). After preprocessing and segmentation, various types of FOMs can be produced that contain observations related to individual cells or regions of interest (ROIs), such as cell neighborhoods or functional tissue units, defined by machine learning algorithms or curated by human experts. Features can include the signal intensity of specific probes or morphological categories such as size and shape. Multidimensional (i.e., > 2 dimensions) FOMs can contain pixel-level intensities per coordinate for each channel in each cell. We note that a robust matrix and analysis metadata standard does not necessarily apply to upstream file types such as raw images, segmentation masks, or sequence alignment and mapping files (SAM/BAM/CRAM). However, having provenance about the tool used for the preprocessing as well as a link to the source file for a derived matrix for a derived FOM is desirable.

### Matrix metadata standards

For each of the major classes of matrix (FOM, FID, OID, FEA, OBS, FNG, ONG, and REC), we develop metadata fields that can be used to describe various aspects of individual matrices (**Table 1**). Primary fields for the FOM class are used to describe the type of biological data being measured (***analyte***), the sets of features and observations that have been included in the matrix (***feature_subset**, **obs_subset***), and the type of processing that was applied to produce the matrix (***processing***). An additional ***modality*** field can be used to denote FOMs containing different data types that require higher levels of integration. In many cases, ***modality*** will be synonymous with ***analyte*** as multimodal workflows seek to cluster cells based on multiple biological modalities. However, ***modality*** is meant to be a broader term that can capture other types of integration as well. For example, researchers may want to integrate the same analyte across different species or across datasets generated with different technologies. The ***processing*** field is used to describe the type of measurement in the FOM from the data analysis perspective. The same biological data may have raw, normalized, and standardized forms. This field can also be used to distinguish reduced dimensional objects such as PCAs and UMAPs from other upstream matrices. One note is that many tools store the biological data matrices and reduced dimensional matrices in separate sections of the data object due to the fact that they are used in different parts of the workflow for different purposes and often contain different features. From the analysis schema perspective, we found that these types of matrices can share similar sets of metadata fields and thus were grouped together into the broader FOM class. The subset fields capture the group of features and cells that belong to the matrix. The ***obs_subset*** field can be used to capture information about the level of filtering that has been applied to the cells or if the cells have been subsetted based on a biological category. Many tools will group all matrices with the same set of observations with the same data object and create separate data objects after applying subsetting operations. This field allows matrices across different data objects within a dataset to be appropriately annotated. Similarly, the ***feature_subset*** field can be used to annotate matrices that contain a subset of detected or highly variable features (e.g. highly variable genes or the PCs explaining the most variance). Other fields such as ***data_type*** and ***representation*** capture some characteristics of the original storage mode of the matrix. These fields are primarily useful when data is converted to simple flat files (e.g. CSV) that do not always have inherent ways of recording this information. More advanced tools and storage formats that have the ability to import flat files can take this into account when converting the FOM to platform-specific data types and matrix representations.

**Table 1.** Description of fields in MAMS.

| Field | Description | Class |
| --- | --- | --- |
| id | Denotes the unique id of the matrix, annotation data frame, or graph and should be unique within the set of objects within the same class of the dataset. | General |
| dataset_id | All matrices and data frames within this group should have observations and features that belong to a superset of observations and features that encompass an entire dataset. | General |
| class | Denotes the class of the matrix, array, data frame. | General |
| data_type | Explicitly describes the type of data stored in the FOM | FOM |
| representation | Preferred representation of the matrix. | FOM |
| obs_unit | Biological unit of the observations | FOM |
| processing | Used to describe the nature of the data contained within the matrix. | FOM |
| analyte | Used to describe the biological analytes being quantified in the matrix. | FOM |
| modality | Describes the modality of the matrix. | FOM |
| obs_subset | Describes the subset of observations that are present in the FOM. | FOM |
| feature_subset | Describes the subset of features that are present in the FOM | FOM |
| parent_id | Denotes the id(s) of the parent matrices that were used to produce the matrix | FOM |
| parent_relationship | Denotes the type of relationship with the parent matrix or matrices | FOM |
| edge_metric | Name of the distance or similarity metric used to create the edges between features. | ONG |
| metric_type | "distance" indicates that smaller values denote more relatedness between features (e.g. euclidean distance) while "similarity" indicates that larger values denote more relatedness between observations (e.g. Pearson correlation) | ONG |
| edge_metric | Name of the distance or similarity metric used to create the edges between features. | FNG |
| metric_type | "distance" indicates that smaller values denote more relatedness between features (e.g. euclidean distance) while "similarity" indicates that larger values denote more relatedness between observations (e.g. Pearson correlation) | FNG |
| record_id | Unique id to denote a combination of entries for record_package_name, record_package_version, record_function_name, and record_function_parameters |  |
| record_package_name | Name of the package, tool, or software that ran the algorithm to produce the matrix, annotation, or graph. | REC |
| record_package_version | Version of the package, tool, or software that ran the algorithm to produce the matrix, annotation, or graph. | REC |
| record_function_name | Name of the function or mathematical operation used to produce the matrix, annotation, or graph. | REC |
| record_function_parameters | Key/value pairs describing the primary parameters and their values used in the function call. | REC |
| record_workflow_link | Public link to workflow that ran the tool. | REC |
| record_runtime_start | The start time of the algorithm or operation. | REC |
| record_runtime_end | The finishing time of the algorithm or operation. | REC |
| record_runtime_duration | The total duration of the algorithm or operation. | REC |

During different steps of an analysis workflow, various operations will create new a FOM from an existing FOM or set of FOMs. In the scRNA-seq example, a full raw count matrix containing droplets can be subsetting to obtain a filtered raw count matrix containing cells and the raw counts in this matrix can be further normalized and log transformed. While the previous metadata fields capture information about the data contained with a FOM, additional metadata fields are needed to capture the relationships between different FOMs. In MAMS, the ***parent_id*** field can be used to link a FOM to one or more parent FOMs and correspond to the arrows in the use cases. The ***parent_relationship*** field defines terms that include different operations to create novel FOMs including transformation, subset, concatenation, reduction, factorization, and aggregation. One particular use case where this information can be useful is for efficient management of concatenated or subsetted FOMs. Creating new FOMs by subsetting or concatenating existing FOMs can create unnecessary copies of existing data and increase storage. However, some data objects are taking advantage of “views” which create a virtual view of a subset of the data without copying the original data^17^. Capturing which matrices are direct subsets or concatenations of other ones in the metadata can further support the use of views across platforms and reduce the overall size of single cell datasets.

Lastly, metadata fields for the other classes were also defined in MAMS. For the ID class, fields are included to denote if an ID is a compound ID separated by a delimiter. The neighborhood graph classes have fields to denote the metric used to quantify the relationship between observations or features as well as a field to denote whether the quantity is a similarity- or distance-based metric (i.e. whether higher or lower values indicate a higher degree of relatedness). The ***dataset_id*** field is a broad term used by all classes to denote a group of related matrices used at any point during an analysis. Lastly, the ***record_id*** is a field used to link matrices or annotations to items in REC class.

### Harmonization of matrix labels using common ontologies

One challenge when merging and harmonizing datasets processed with different workflows is that the same type of FOM may be given different labels. For example, a normalized matrix may be called “data” “normcounts”, or given any label by the user running the workflow depending on the tool. For several of the metadata fields, we developed a set of harmonized ontologies for commonly generated matrices. Example terms for the ***processing*** field is shown in **Table 2**. For ***processing***, “raw” denotes a general term for the original measurements derived directly from the source files. The term “counts” denotes raw measurements that are integers such as UMI-corrected read counts for mRNA, protein, or ATAC-seq data. The term intensity denotes raw measurements that are derived from fluorescent intensities commonly used in imaging-based techniques. Other terms such as “normalized”, “lognormalized”, and “scaled” can be used to describe the stage of data processing on the original features. The “reduction” term denotes reduced dimensional representations used for input into downstream analysis (e.g. PCA) while the “embedding” term is reserved for low dimensional representations often used for visualization (e.g. UMAP). The ***analyte*** field has terms such as rna, protein, chromatin, dna, lipid, metabolite, and morphology to describe the biological feature captured by the measurements in the FOM. Terms for the ***obs_subset*** field include “full” to denote a FOM that has all original observations, “filtered” to denote observations that enough total signal above background. (e.g. true cells in droplet-based scRNA-seq), “detected” to denote observations that have minimum levels of detection across features, “nonartifact” which can be used as a general term to describe filtering that may occur due other quality control metrics, and “clean” to denote an “analysis ready” set of observations.

The majority of fields also have corresponding description fields used to describe the selected term. We have also supplied descriptions for the suggested ontologies (see Table 2 for an example with *processing_description*). While some ontological terms are suggested in MAMS, any text can be supplied to allow the metadata fields to adapt to future scenarios. Thus, researchers have the flexibility to use and define custom terms and descriptions that do not fit the current set of suggested ontologies. Using a set of predefined ontologies with flexibility to add new terms will promote harmonization across platforms and technologies while having the flexibility to adapt to novel analysis workflows.

**Table 2.** Suggested categories for the *processing* and *processing_description* field with examples.

| processing | processing_description | Notes/Examples |
| --- | --- | --- |
| raw | Original measurements have not been altered |  |
| counts | Raw data for assays that produce integer-like data such as scRNA-seq. Child of `raw`. |  |
| intensities | Raw data for assays that produce continuous data. Child of `raw`. |  |
| decontaminated | Measurements have been corrected for background signal. | Ambient RNA removal in single-cell RNA-seq. |
| lograw | The log of the raw data |  |
| logcounts | The log of the raw counts |  |
| logintensities | The log of the raw intensity values. |  |
| corrected | Measurements have been corrected for observation-level covariates |  |
| normalized | Data that has been normalized for differences in overall signal abundance between observations. | Correcting for total number of UMIs or reads in each cell. |
| lognormalized | Data that has been log transformed after normalizing for differences in overall signal abundance between observations. Child of `normalized`. |  |
| centered | Data with features have been made to center around a standard quantity such as the mean or median. | Mean-centered data |
| scaled | Data with features have been centered around a standard quantity and standardized to have similar variances or ranges. | Z-scored data |
| reduction | A matrix containing a data dimensionality reduction generally useful for input into tools for downstream analysis such as clustering or 2D-embedding. | PCA, ICA, Autoencoders |
| embedding | A matrix containing a low dimensional embedding (usually 2D or 3D) generally used for visualization. Child of `reduction` | UMAP, tSNE |

### Provenance related metadata for of analysis of matrices

The FAIR Data Principles (Findable, Accessible, Interoperable, and Reusable) are a set of guiding principles to support the reusability of data^22^. A major component of reproducibility is that the data should have information related to provenance including how it was generated, preprocessed, and analyzed. A major goal of many groups has been to develop metadata standards related to the demographics of the donor or patient include disease relevant phenotypes as well as information about the technologies and protocols used to create data from biological specimens. However, once the raw data has been generated, the degree to which the details of the software and methods that create new FOMs and annotations is variable and limited across groups. Some data processing centers will have a central workflow for processing which can be accessed via a repository. Some information about the software and analysis parameters may also be listed in the publication. However, this information is not standardized and might not be stored along with the FOMs in the file format being used.

The ***record_package_name*** describes the name of the package, tool, or software that ran the algorithm to produce the matrix, annotation, or graph. The ***record_package_version*** denotes the version of the package, tool, or software that ran the algorithm to produce the matrix, annotation, or graph. The ***record_function_name*** describes the name of the function or mathematical operation used to produce the matrix, annotation, or graph. The ***record_function_parameters*** is a list containing key-value pairs describing the primary parameters and their values used in the function call. Finally, the ***record_workflow_link*** can be used to denote a public link to a repository containing the workflow that ran the tool. For example, links to workflow scripts encoded in CWL, WDL, Nextflow, Snakemake are often stored in a public repository such as GitHub or DockerHub^23–25^. In summary, these fields can be used to record provenance information about matrices or individual annotations and will aide in reproducibility of single-cell data

### Implementation of MAMS in a portable format

In order to facilitate the adoption of MAMS, we developed a simple list-like structure that can be used to record MAMS metadata fields for matrices in a dataset. This structure can be stored in configuration file formats such as JSON and YAML and can record information about the dataset even if the FOMs are stored across different data objects or file formats. Each class of matrix has an entry within the list for each dataset and each matrix is denoted with a unique ID. The MAMS fields are then categorized as key-value pairs under each individual matrix. Several additional fields are implemented within this format to describe the location of each matrix and potential relationship between matrices. The ***filepath*** field is used to describe the path to the file or data object containing the matrix which can point to a flat file, HDF5 object, or other programming language specific storage format (e.g. rds file format for R). The ***accessor*** field is used to denote the command to retrieve the matrix from a data object. An additional set of linking fields (***oid, fid, obs, fea***) can be used to capture the relationships between FOMs and annotation or IDs matrices. Using these linking fields allow potential relationships between FOMs and other matrix classes to be maintained independently of file path or memory location within a data object. This format can serve as an intermediate standard to store MAMS information even if the underlying data object does not have the capability to store this type of metadata.

## DISCUSSION

Metadata related to analysis and provenance of FOMs is important for the reanalysis and reproducibility of single-cell data but has been inconsistently curated and captured across software platforms and data coordinating centers. In order to facilitate the harmonization of metadata standards related to data analysis across groups, we created MAMS with input from multiple consortia and software development groups. Many different combinations of tools and parameters can be applied during an analysis workflow to produce different numbers and types of matrices. By curating analysis “use cases” from several existing workflows involving multiple data types and analysis goals, we have characterized several core principles that can be used to annotate data matrices. The ability of MAMS to capture these principles will allow for the curation of matrices generated from diverse settings and support future complex workflows.

One of the major challenges when integrating matrices from different datasets is to determine which matrices should be selected and merged. Having fully annotated matrices with MAMS fields can aid in making this process more systematic. For example, having fields to clearly denote ***analyte*** and ***modality*** can ensure that matrices capturing similar biological measurements will be appropriately merged. Various integrative efforts may want to use different data matrices depending on the goals of the analysis. Reprocessing efforts will likely want to start with the matrices containing the most raw and unfiltered form of the data while other efforts seeking to answer targeted biological questions may just want to merge the filtered and normalized matrices for each dataset. Overall, better metadata standards such as MAMS are becoming increasingly important with the goal of re-analysis and integration across datasets.

Different workflows and tools may not save every matrix produced during an entire workflow. For example, PCA matrix may be calculated from a matrix of highly variable scaled genes that is not permanently stored within the workflow. The goal of MAMS is not to require that every matrix produced by an analysis workflow should be stored in a file or data object or maintained in a repository. Rather the goal of MAMS is to ensure that every matrix stored within a data object or file format can be properly annotated with metadata fields and relevant provenance records.

In general, the metadata fields defined by MAMS are not dependent on specific formats or data storage standards and can be implemented in any existing software that organizes matrices under a common API. However, updating existing software packages can take substantial time and effort. Therefore, we developed a platform-agnostic list-like structure to store MAMS fields that can be stored in YAML or JSON formats. This file can serve as a “dataset configuration” file which can be used in conjunction with any underlying storage formats. Currently, the curation of these attributes is not automated in most analysis workflows and software packages. Future efforts will be needed to implement these standards across platforms and relieve downstream users of the task of manual curation. Overall, the successful implementation of these data analysis metadata standards will facilitate harmonization and integration of datasets stored across different platforms or repositories and produce single-cell data that more closely aligns with FAIR standards.

## METHODS

We assembled a working group consisting of members from various data curation centers, software platforms, academic and industrial institutions. The working group met monthly over the course of a year to discuss various aspects of data curation and file formats. “Use cases” for analysis of single-cell data were derived from Seurat^14,15,26^, Scanpy^19^, and Bioconductor example workflows^16^. Imaging-based workflows were based on a workflow from MCMICRO^27^. Metadata fields were defined based on matrices and provenance fields produced by these workflows or based on use cases from the experience of the working group members. Multi-modal PBMC data was accessed from https://www.10xgenomics.com/resources/datasets/pbm-cs-of-a-healthy-donor-5-gene-expression-and-cell-surface-protein-1-standard-3-0-0. MAMS documentation, use cases, and example files can be found at https://github.com/single-cell-data/mams/.

## ACKNOWLEDGEMENTS

This work was funded by the NCI Human Tumor Atlas Network (HTAN) Lung Pre-Cancer Atlas (PCA) 1U2CCA233238-01 (J.D.C.); Single Cell Gene Expression Atlas grant from the Wellcome Trust 108437/Z/15/Z (P.M.); National Heart, Lung, and Blood Institute grant U24HL148865 (D.J.S.).

## AUTHOR CONTRIBUTIONS

Methodology (Y.W, I.S., A.S., R.D., B.R.H., H.H.C., A.M., D.J.S., J.H., J.D.C.), Data Curation (Y.W., W.K.T.), Funding acquisition (N.G., J.D.C.), Conceptualization (J.D.C., K.D., N.G., T.T.), Writing - Original Draft (Y.W, J.D.C.), Writing - Review & Editing (A.S., A.M., D.J.S., P.M.).

## DECLARATIONS

The authors declare that they have no competing interests.

## SUPPLEMENTARY FIGURES

**Supplementary Figure 1.**
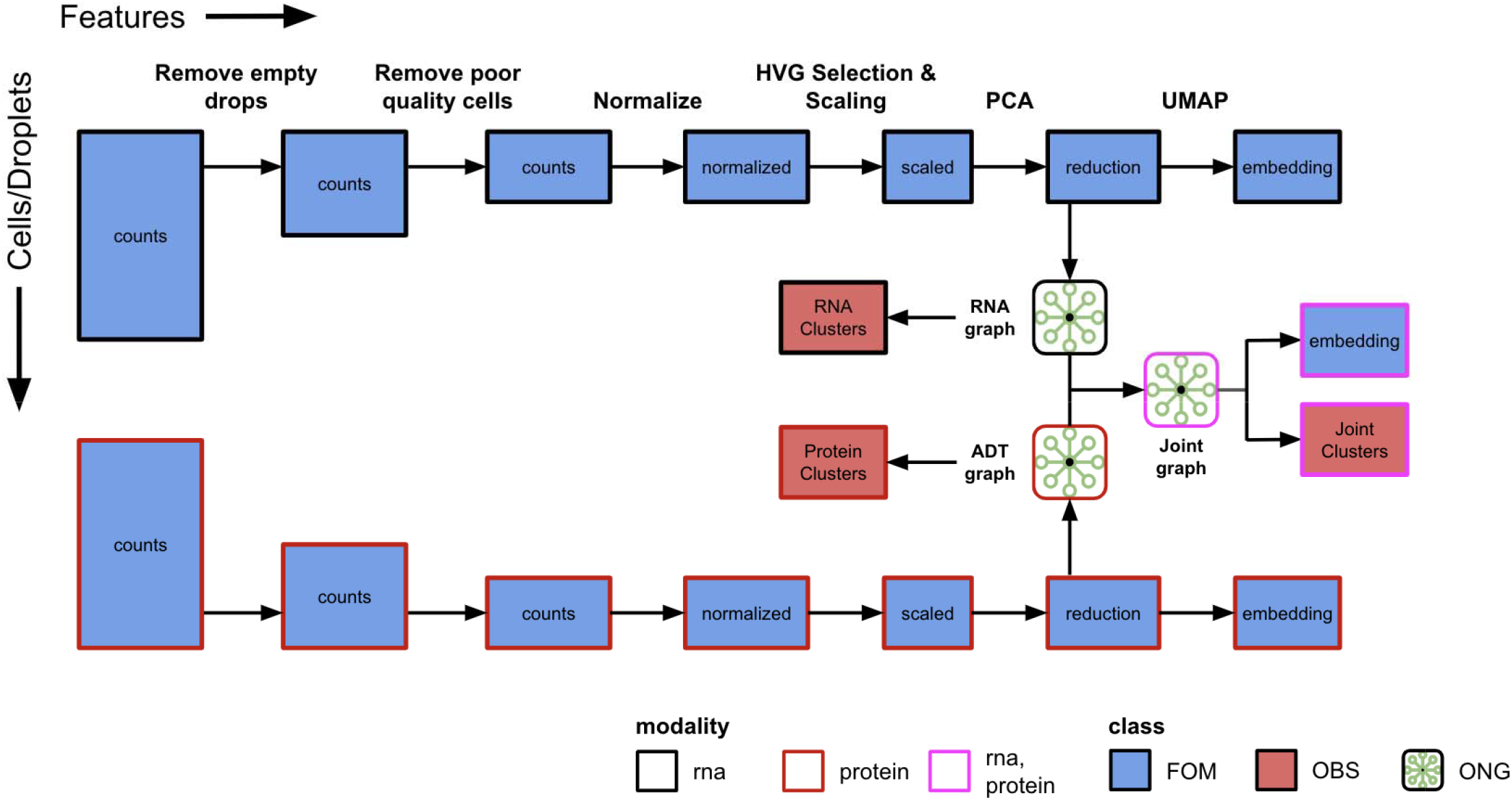
Multi-modal single cell RNA-seq and CITE-seq analysis workflow. Several technologies can produce multi-modal data by capturing different biological analytes on the same sets of observations. This workflow demonstrates FOMs and annotations for single cells that are produced by joint profiling of RNA and protein expression using CITE-seq. A separate data matrix is generated for each modality (box color) and will undergo filtering for empty drops and poor-quality cells. Although the same set of cells are retained for analysis in this illustration, different sets of filtering criteria may be applied to each modality independently. Log normalization, scaling, dimensionality reduction, 2-D embedding, and clustering can be performed in parallel for each modality. Further analysis can be performed using a nearest neighbor graph generated by combining the individual neighbor graphs from each modality. A new 2-D embedding and set of cluster labels can be generated with both underlying modalities. The “modality” field can be a list to denote the combination of analytes that were used in the generation of the FOM. Feature annotation and ID classes are not shown.

**Supplementary Figure 2.**
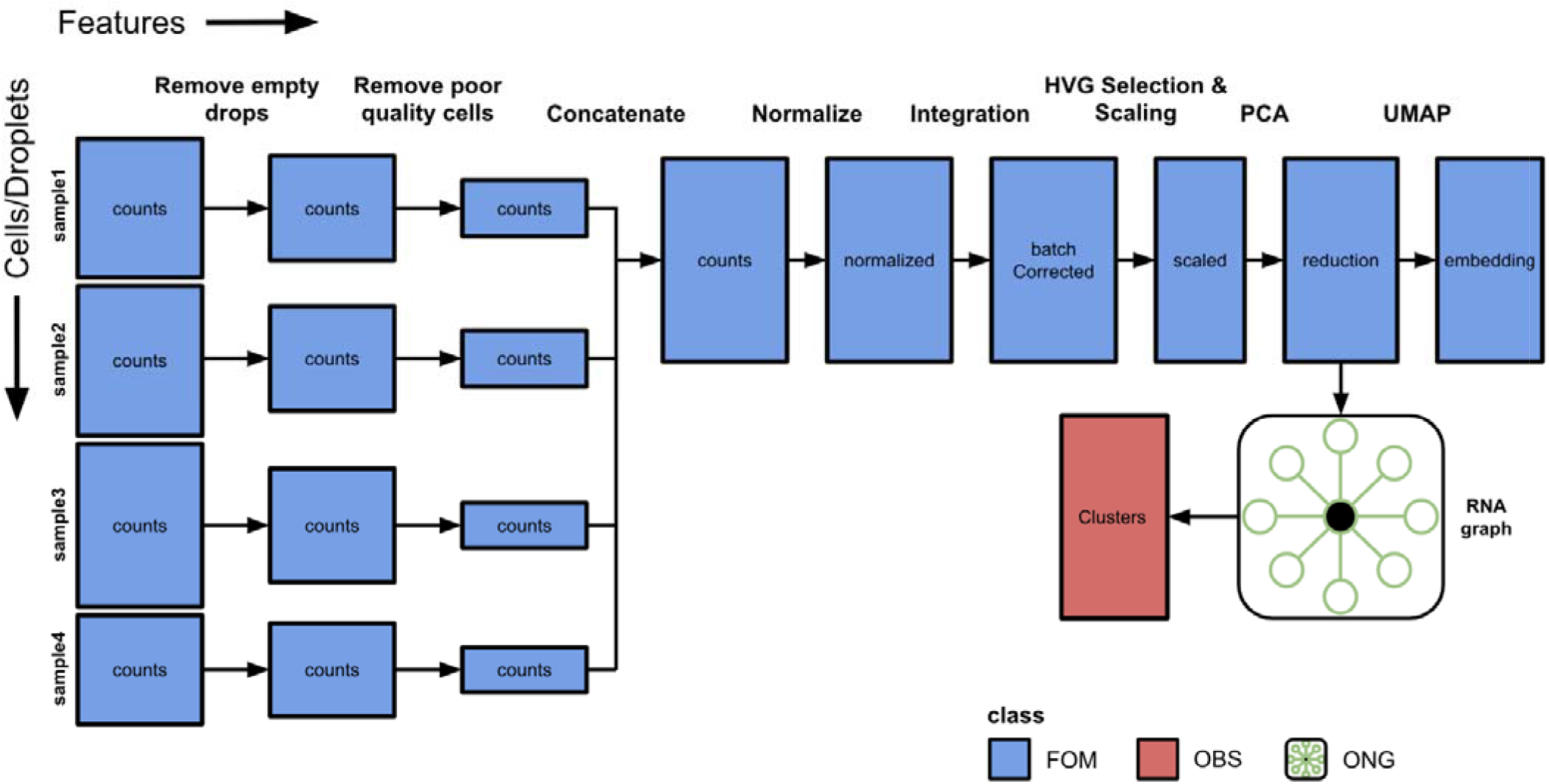
Multi-sample single-cell analysis workflow with batch correction. In many datasets, data will be generated for multiple samples and jointly analyzed. This workflow demonstrates data matrices from four samples with different numbers of observations. Empty droplets are identified and removed from each sample individually followed by removal of poor quality cells or artifacts such as doublets/multiplets. Filtered matrices from all samples are then concatenated to produce a combined cell matrix. Note that the order of filtering may be different in various workflows (e.g. filtering of poor quality cells may occur after concatenation). The combined matrix of all samples can be analyzed using standard workflows for normalization, scaling, dimensionality reduction, 2-D embedding, and clustering. In some circumstances, technical differences between samples could produce unwanted clustering. Integration and batch correction methods can be applied at different steps depending on the algorithm or statistical method. In this example, a batch correction algorithm is applied to the log normalized data to produce a new matrix which can be used in subsequent steps. Other variations of this workflow may include using a batch correction or integration method that works on the reduced dimensional matrices to produce a new reduced dimensional object which can subsequently be used to generate a graph.

**Supplementary Figure 3.**
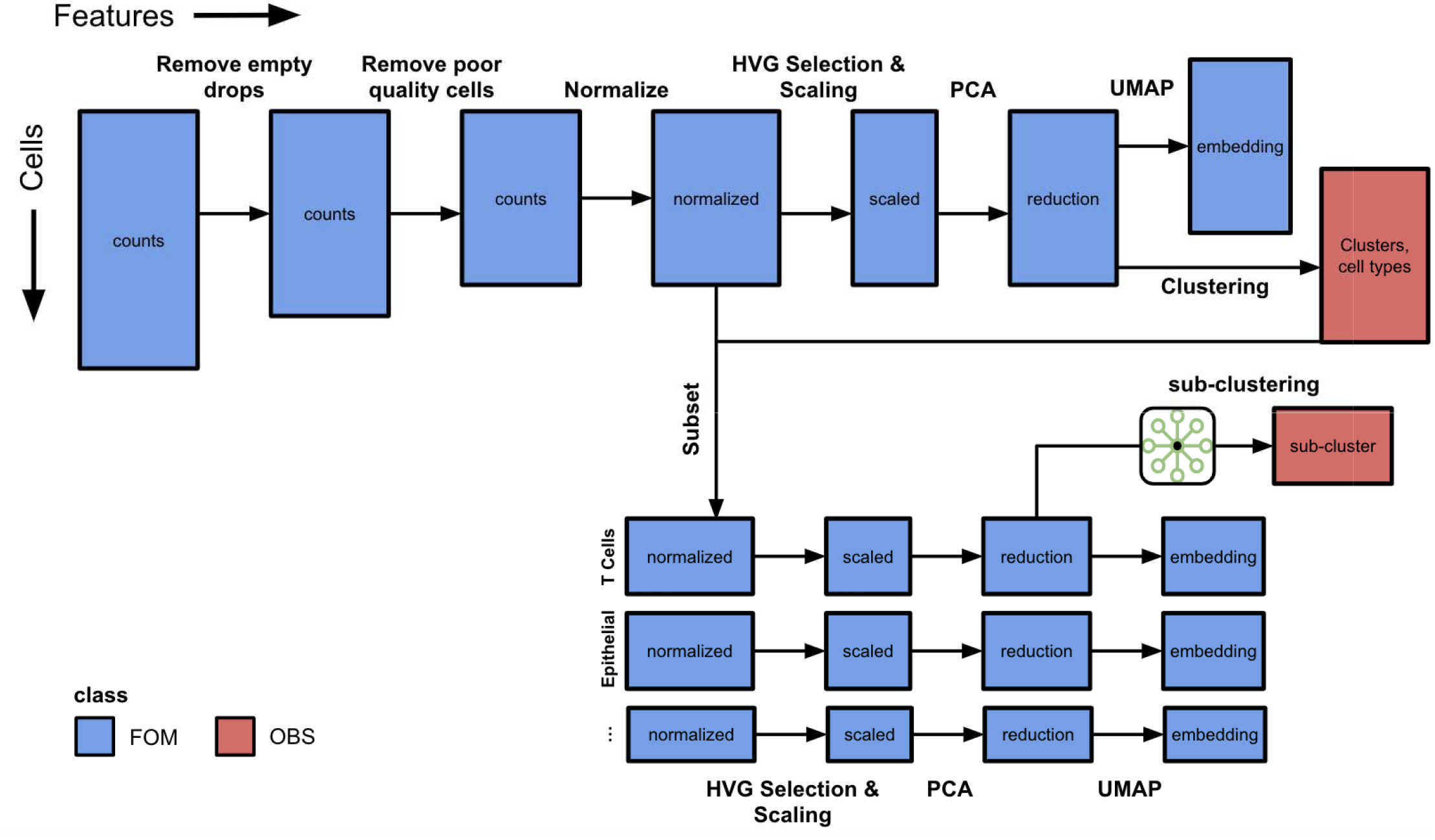
Single-cell RNA-seq workflow with additional analysis of biological subsets. During the analysis of real-world datasets, subsets of data are obtained for additional downstream analysis. In this example, a matrix of scRNA-seq data is taken through the standard analysis workflow including filtering, normalization, scaling, dimensionality reduction, and clustering. While this workflow may identify the major cell types present in the dataset, further clustering of individual cell types may reveal additional heterogeneity. The normalized matrix is subsetted using cell type labels derived from clustering. The *obs_subset* field can be used to denote biological subsets of observations such as T-cell, epithelial cell, etc. Each subsetted normalized matrix will undergo a similar workflow of scaling, dimensionality reduction, 2-D embedding, and clustering to identify subpopulations within each major cell type. Feature annotation and ID classes are not shown.

**Supplementary Figure 4.**
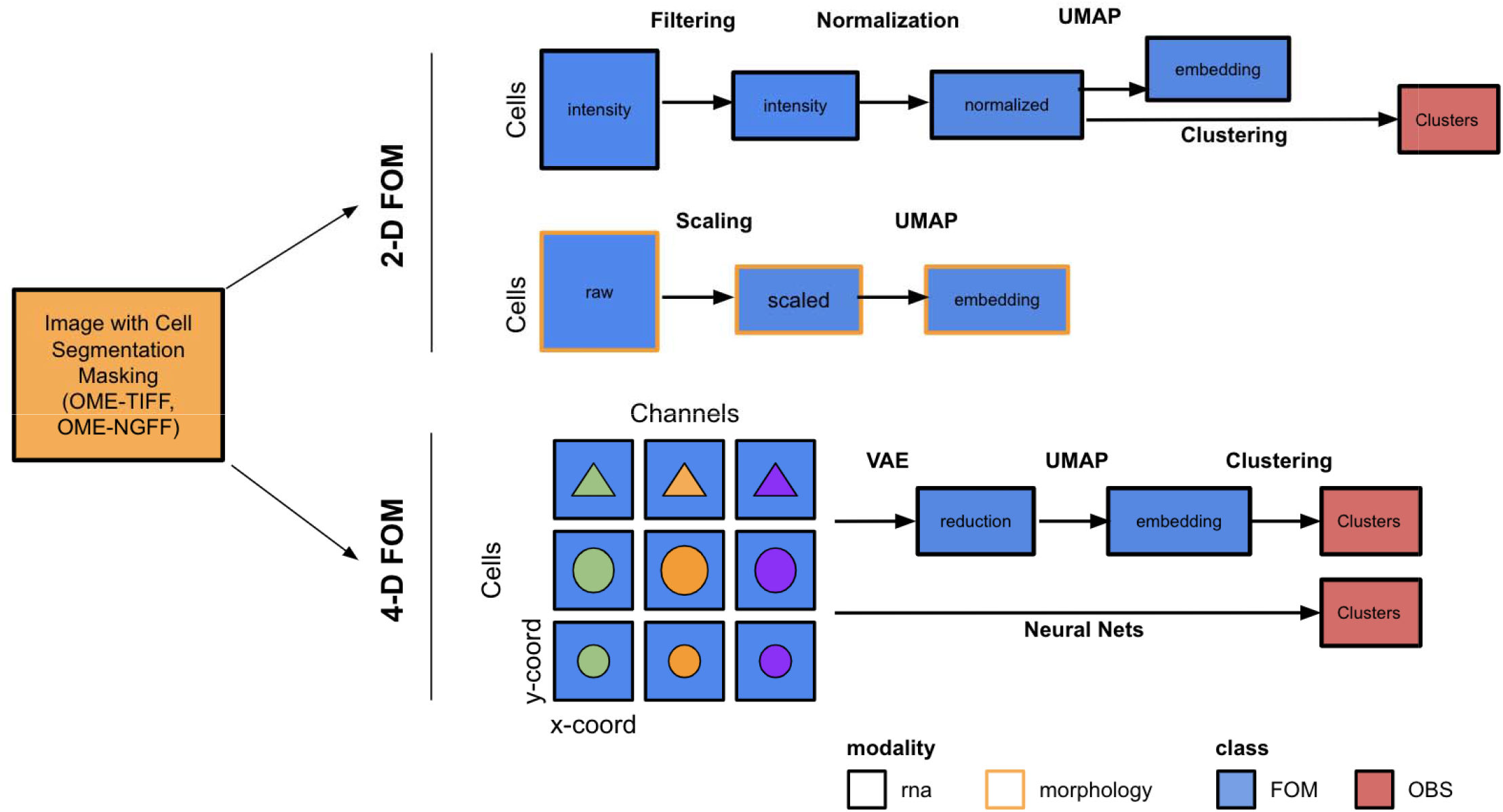
Imaging-based analysis workflow that produces FOMs. Several imaging-based technologies can examine tissue slides to quantify molecular and cellular features in 2-D or 3-D space. Image analysis methods (e.g., segmentation) or manual curation by an expert can be used to identify Regions of Interest (ROIs) such as cell neighborhoods on each slide. Spatial technologies yield two flavors of FOMs. First, 2-dimensional FOMs can be derived that quantify the intensity levels of each marker in each cell or ROI. These raw intensity matrices can be augmented by additional morphological characteristics, such as cell size and shape, as well as spatial motifs within ROIs. The 2D FOMs can be filtered, normalized and used in clustering and embedding workflows in a similar fashion to single-cell sequencing FOMs. Second, a 4-D FOM can be generated that stores the pixel-level intensities along the x- and y-coordinates in each cell or ROI for each channel/marker. This type of FOM can be analyzed with algorithms that work directly on pixel-level data, such as variational autoencoders, to produce a reduced dimensional matrix which can be used in clustering and embedding workflows. Feature annotation and ID classes are not shown.

## REFERENCES

1. Regev, A. et al. The Human Cell Atlas. Elife 6, (2017).

2. HuBMAP Consortium. The human body at cellular resolution: the NIH Human Biomolecular Atlas Program. Nature 574, 187–192 (2019).

3. Rozenblatt-Rosen, O. et al. The Human Tumor Atlas Network: Charting Tumor Transitions across Space and Time at Single-Cell Resolution. Cell 181, 236–249 (2020).

4. Li, H. et al. Fly Cell Atlas: A single-nucleus transcriptomic atlas of the adult fruit fly. Science 375, eabk2432 (2022).

5. Plant Cell Atlas Consortium et al. Vision, challenges and opportunities for a Plant Cell Atlas. Elife 10, (2021).

6. Gaddis, N. et al. LungMAP Portal Ecosystem: Systems-Level Exploration of the Lung. Am J Respir Cell Mol Biol (2022) doi:10.1165/rcmb.2022-0165OC.

7. Ardini-Poleske, M. E. et al. LungMAP: The Molecular Atlas of Lung Development Program. Am J Physiol Lung Cell Mol Physiol 313, L733–L740 (2017).

8. Clough, E. & Barrett, T. The Gene Expression Omnibus Database. Methods Mol Biol 1418, 93–110 (2016).

9. Sarkans, U. et al. From ArrayExpress to BioStudies. Nucleic Acids Res 49, D1502–D1506 (2021).

10. Puntambekar, S., Hesselberth, J. R., Riemondy, K. A. & Fu, R. Cell-level metadata are indispensable for documenting single-cell sequencing datasets. PLoS Biol 19, e3001077 (2021).

11. Bolewski, J. & Papadopoulos, S. Managing massive multi-dimensional array data with TileDB: — Invited demo paper. in 2017 IEEE International Conference on Big Data (Big Data) 3175–3176 (2017). doi:10.1109/BigData.2017.8258296.

12. Virshup, I., Rybakov, S., Theis, F. J., Angerer, P. & Wolf, F. A. anndata: Annotated data. bioRxiv 2021.12.16.473007 (2021) doi:10.1101/2021.12.16.473007.

13. Bredikhin, D., Kats, I. & Stegle, O. MUON: multimodal omics analysis framework. Genome Biol 23, 42 (2022).

14. Butler, A., Hoffman, P., Smibert, P., Papalexi, E. & Satija, R. Integrating single-cell transcriptomic data across different conditions, technologies, and species. Nat Biotechnol 36, 411–420 (2018).

15. Stuart, T. et al. Comprehensive Integration of Single-Cell Data. Cell 177, 1888–1902.e21 (2019).

16. Amezquita, R. A. et al. Orchestrating single-cell analysis with Bioconductor. Nat Methods 17, 137–145 (2020).

17. Sarfraz, I., Asif, M. & Campbell, J. D. ExperimentSubset: An R package to manage subsets of Bioconductor Experiment objects. Bioinformatics (2021) doi:10.1093/bioinformatics/btab179.

18. Ramos, M. et al. Software for the Integration of Multiomics Experiments in Bioconductor. Cancer Res 77, e39–e42 (2017).

19. Wolf, F. A., Angerer, P. & Theis, F. J. SCANPY: large-scale single-cell gene expression data analysis. Genome Biol 19, 15 (2018).

20. Stuart, T. & Satija, R. Integrative single-cell analysis. Nat Rev Genet 20, 257–272 (2019).

21. Adossa, N., Khan, S., Rytkönen, K. T. & Elo, L. L. Computational strategies for single-cell multi-omics integration. Comput Struct Biotechnol J 19, 2588–2596 (2021).

22. Wilkinson, M. D. et al. The FAIR Guiding Principles for scientific data management and stewardship. Sci Data 3, 160018 (2016).

23. di Tommaso, P. et al. Nextflow enables reproducible computational workflows. Nat Biotechnol 35, 316–319 (2017).

24. Ahmed, A. E. et al. Design considerations for workflow management systems use in production genomics research and the clinic. Sci Rep 11, 21680 (2021).

25. Mölder, F. et al. Sustainable data analysis with Snakemake. F1000Res 10, 33 (2021).

26. Satija, R., Farrell, J. a, Gennert, D., Schier, A. F. & Regev, A. Spatial reconstruction of single-cell gene expression data. Nat Biotechnol (2015) doi:10.1038/nbt.3192.

27. Schapiro, D. et al. MCMICRO: a scalable, modular image-processing pipeline for multiplexed tissue imaging. Nat Methods 19, 311–315

